# Bridging Language Markers and Pathology: Correlations Between Digital Speech Measures and Surrogate CSF Biomarkers in Alzheimer’s Disease

**DOI:** 10.1101/2025.11.05.686472

**Authors:** Yijiang Pang, Liu Chen, Hiroko H. Dodge, Jiayu Zhou

## Abstract

**Background:** Digital language markers show promise in detecting early cognitive impairment related to Alzheimer’s disease (AD), yet their relationship with cerebrospinal fluid (CSF) biomarkers of AD pathology remains unclear mainly due to the lack of data with both CSF and language markers.

**Objective:** This study aims to build links between digital language markers and fluid biomarkers through surrogate CSF biomarkers.

**Methods:** Using NACC clinical data as anchor variables, language makers in the I-CONECT study were linked to NACC CSF data. Surrogate CSF biomarkers were created for I-CONECT subjects using machine learning models from common NACC clinical variables. Correlations assessed associations between CSF and language markers.

**Results:** Lower predicted amyloid-β correlated significantly with reduced syntactic complexity and shorter speech responses. Higher predicted total tau and phosphorylated tau correlated with reduced syntactic complexity.

**Conclusions:** This study demonstrates novel links between language markers and fluid biomarkers, highlighting conversational language as a potential accessible, non-invasive approach for early detection and monitoring of Alzheimer’s pathology.

## Introduction

Alzheimer’s disease (AD) is characterized by progressive cognitive decline associated with amyloid-beta (Aβ) plaques and tau neurofibrillary tangles in the brain. These pathological hallmarks can be reliably measured through cerebrospinal fluid (CSF) biomarkers, notably Aβ1–42, phosphorylated tau (e.g., P-tau181, P-tau217), and total tau (T-tau). While considered gold-standard indicators of AD neuropathology, obtaining CSF biomarkers typically requires invasive lumbar punctures, creating significant barriers to widespread clinical application. Consequently, research interest has grown toward identifying non-invasive, cost-effective biomarkers that could serve as proxies for AD pathology.

Recently, language characteristics have emerged as promising digital biomarkers for detecting early cognitive changes associated with mild cognitive impairment (MCI) and early AD stages [1]. Computational language analysis has demonstrated potential in distinguishing cognitively impaired individuals from healthy older adults, utilizing features such as lexical diversity, syntactic complexity, and psycholinguistic content measured by tools like Linguistic Inquiry and Word Count (LIWC). Over the past decade, literature has shown that the way people speak and use language tends to change in prodromal and early AD, even before overt dementia [1]. More recently, natural language processing (NLP) and machine learning have enabled fine-grained analysis of speech transcripts to detect these subtle linguistic effects of cognitive decline [2–5].

While speech and language biomarkers have shown strong predictive performance for cognitive impairment, measuring their direct connection to AD pathology remains a challenge. This is an important gap, because AD pathology can accumulate silently for years before clinical dementia – some individuals with substantial amyloid/tau burden are cognitively normal, and conversely some cognitively impaired patients have non-AD pathologies. Understanding if and how language characteristics relate to underlying amyloid and tau would validate digital language markers as indicators of disease biology, not merely proxies for clinical status. To date, most digital speech studies have focused on predicting clinical diagnoses (e.g., MCI vs. normal aging) or cognitive test scores [2–5], rather than directly measuring AD neuropathology. There are some pioneering studies that have explored the connection between speech/language and AD fluid biomarkers [6–10], providing concrete evidence of the linking between digital speech markers with CSF biomarkers of AD pathology. However, these studies are constrained to relatively small samples where both modalities are available, and the highly structured speech tasks overlook comparability and standardization in speech data collection. One major barrier to establishing these associations is the practical difficulty of collecting simultaneous speech and fluid biomarker data within the same cohort. The Internet⍰Based Conversational Engagement Clinical Trial (I⍰CONECT) [6] provides a rich dataset of semi⍰structured conversation for analyzing language patterns associated with MCI. However, the I-CONECT participants did not undergo CSF biomarker assays as part of the trial. To overcome this limitation, we developed a surrogate biomarker modeling strategy: using the expansive National Alzheimer’s Coordinating Center (NACC) database, we trained machine learning models to predict CSF Aβ42, P-tau181, and T-tau levels from a broad array of non-invasive measures. Using shared clinical variables available in both the NACC dataset and the I-CONECT trial, we trained machine learning models to predict CSF biomarker levels in NACC participants. These validated predictive models were subsequently applied to estimate surrogate CSF biomarker values for I-CONECT participants, who lacked fluid biomarker measures. This approach uniquely enables the assessment of correlations between surrogate CSF biomarkers of AD pathology and naturalistic speech and language markers within a socially engaged, conversational cohort.

Given the lack of the comprehensive datasets that jointly capture semi-structured spoken language data and fluid biomarker measures, this study is, to our knowledge, the first systematic investigation linking digital speech markers with surrogate CSF biomarkers of AD pathology in a naturalistic conversational context, which demonstrates the feasibility of performing scalable analysis from separate speech and biomarker cohorts. This approach exemplifies the transformative potential of integrating traditional biological biomarkers with innovative digital health technologies, leveraging decades of accumulated clinical data from Alzheimer’s Disease Research Centers (ADRCs) and NACC to pave new pathways for early, accessible, and non-invasive detection and monitoring of Alzheimer’s disease.

## Methods

### Datasets and Common Variables

We leveraged two primary datasets: the I-CONECT trial and the NACC database. I-CONECT (NCT02871921) is a multi-site randomized controlled trial that delivered frequent, semi-structured conversational engagements via webcam to socially isolated older adults (age ≥75) with normal cognition or mild cognitive impairment (MCI) [11]. Over 6–12 months of intervention, I-CONECT collected rich language data from participants’ spoken conversations, along with standard cognitive and health assessments using NACC forms. Briefly, participants engaged in 30⍰minute, semi⍰structured video⍰chat conversations up to four times per week for six months (induction), followed by twice⍰weekly sessions for another six months (maintenance). Interviewers and participants discussed a daily topic with accompanying picture prompts in a natural conversational setting. These recordings have been widely used to assess the ability of linguistic and acoustic features to distinguish mild cognitive impairment (MCI) from those with normal cognition [4,12–14]. The NACC (National Alzheimer’s Coordinating Center) database provided a comparative clinical dataset drawn from ADRCs across the US [15]. NACC, established in 1999, contains extensive uniform clinical evaluations and biomarker data from ADRC participants, serving as a cumulative research repository. Crucially for this study, a subset of NACC participants have measured CSF biomarker values for amyloid-β 1–42 (Aβ42), total tau (T-tau), and phosphorylated tau 181 (P-tau181) [16]– the well-established CSF markers of AD pathology [17]; see Table 1 for demographic and characteristics regarding each biomarker.

**Table 1.** Demographic and clinical characteristics of participants from the I-CONECT and NACC cohorts, including age, sex, education, race, MoCA scores, and cognitive status for participants.

| NACC dataset |  |  |  |  |
| --- | --- | --- | --- | --- |
| Variables | | Aβ42 | P-tau181 | T-tau |
| Number of Unique Subjects |  | 2055 | 1875 | 1986 |
| Sex: Female, % |  | 50.9 | 50.1 | 50.7 |
| Age, years | | 70.5±8.9 | 70.3±9.0 | 70.5±8.9 |
| Education, years | | 16.0±5.7 | 15.9±6.0 | 16.0±5.8 |
| Race (%) | White | 89.9 | 89.5 | 89.6 |
|  | Black or African American | 6.3 | 6.9 | 6.5 |
|  | American Indian or Alaska Native | 0.1 | 0.1 | 0.1 |
|  | Native Hawaiian or Other Pacific Islander | 0 | 0.1 | 0.1 |
|  | Asian | 1.8 | 1.1 | 1.7 |
|  | Other or Unknown | 2.0 | 2.2 | 2.1 |
| MoCA score |  | 23.2±5.8 | 22.8±6.0 | 23.1±5.9 |
| Cognitive Status (%) | Normal | 51.7 | 51.6 | 51.6 |
|  | MCI | 29.8 | 29.5 | 29.6 |
|  | AD | 15.2 | 15.6 | 15.6 |
| **I-CONNECT dataset** |  |  |  |  |
| Number of Unique Subjects |  | 186 |  |  |
| Sex: Female, % |  | 69.9 |  |  |
| Age, years |  | 81.1±4.5 |  |  |
| Education, years |  | 15.2±2.2 |  |  |
| Race (%) | White | 80.1 |  |  |
|  | Black or African American | 18.8 |  |  |
|  | American Indian or Alaska Native | 0 |  |  |
|  | Native Hawaiian or Other Pacific Islander | 0 |  |  |
|  | Asian | 0.5 |  |  |
|  | Other or Unknown | 0.5 |  |  |
| MoCA score |  | 23.5±3.5 |  |  |
| Cognitive Status (%) | Normal | 56.5 |  |  |
|  | MCI | 43.5 |  |  |

We identified 168 common variables collected in both datasets, which include demographics (age, education, etc.), physical (weight, resting heart rate, etc.), cognitive test scores (memory recall, fluency, etc.), functional ratings, and other clinical measures. A table describing the shared variables is provided in Appendix: Supplementary Table 1. These overlapping features enabled a harmonized feature space between NACC and I-CONECT, laying the groundwork for our surrogate biomarker modeling.

### Surrogate CSF Biomarker

Our approach used the NACC data as a training set to learn mappings from clinical features to CSF biomarker values, and then applied these models to I-CONECT participants who lack actual CSF measures. Workflow of the proposed method is demonstrated in Figure 1.

**Figure 1.**
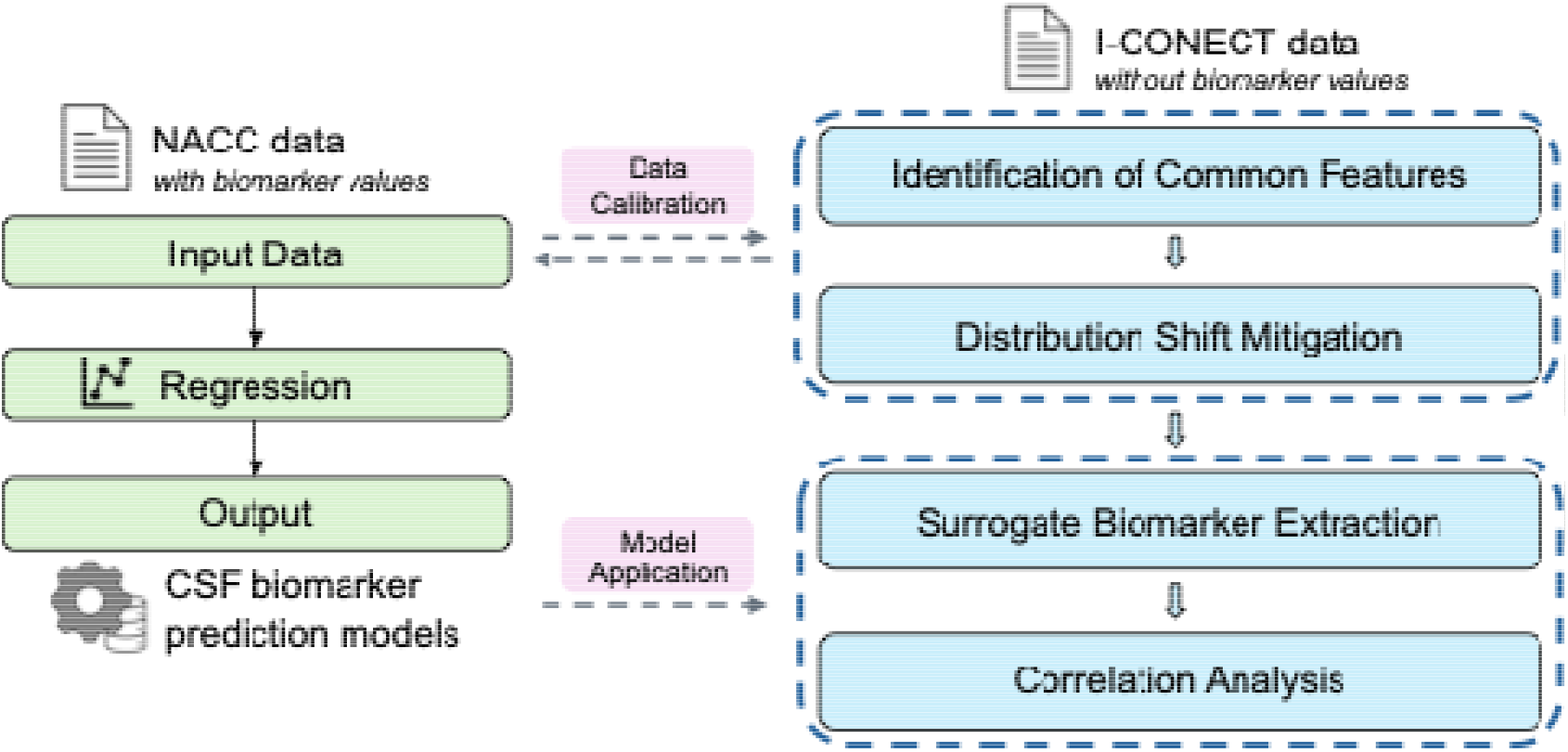
shows the workflow of the proposed method. CSF biomarker prediction models trained exclusively using NACC data; interacting with the NACC data and prediction models, I-CONECT participants are further assigned surrogate biomarker labels, enabling correction analysis with linguistic features.

We trained separate regression models for each CSF biomarker (Aβ42, P-tau181, T-tau) using LightGBM (Light Gradient Boosting Machine), a gradient-boosting decision tree framework known for efficiency and accuracy with high-dimensional data [18]. Model development followed a rigorous procedure: we split the NACC sample with CSF data into 90% for training (with 5-fold cross-validation for hyperparameter tuning) and 10% held-out for testing. The models learned to predict continuous CSF values from the 168 features. Predictive performance was evaluated by cross-validated R^2^ and mean squared error on the training folds and by examining error distributions on the hold-out set. Once obtaining the CSF biomarker prediction models, we next predicted subjects’ CSF biomarkers in I-CONECT through the common variables. Particularly, a step aiming to mitigate the potential distribution shift between NACC and I-CONECT is employed and detailed below.

### Distribution Shift Assessment

As shown in Table 1, we observed notable differences in sex and age between the two datasets, with potential variations in other common variables. Given that machine learning models are generally expected to perform well on in-distribution testing data, we assessed whether the I-CONECT and NACC datasets were drawn from similar distributions in the 168-dimensional feature space, aiming to identify and mitigate potential out-of-distribution data points. A Principal Component Analysis (PCA) of the combined dataset (marking data source) revealed a distribution shift: I-CONECT participants clustered somewhat separately from NACC participants along the first few principal components. This reflects differences in the two cohorts (e.g., I-CONECT’s socially isolated older adults recruited from communities vs. NACC’s clinic-based cohort). A few I-CONECT cases lay at the fringes of the NACC data cloud, indicating they had feature combinations not well-represented in the NACC training set. To mitigate potential extrapolation error, we performed an outlier trimming on I-CONECT data before prediction: specifically, we identified individuals with extreme PCA scores outside the range of the NACC sample and removed these outliers in a subsequent sensitivity analysis.

### Feature Importance of Predictive Models

To interpret the CSF biomarker prediction models, we analyzed the feature importances, specifically gain importance, for each LightGBM model. The gain importance directly measures the reduction in impurity achieved by a feature across all splits where that feature is used in the decision trees, which is formulated as: 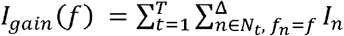 where *T* is the total number of trees in the model, *N*_*t*_ is the set of all nodes in the *t*-*th* tree, *f*_*n*_ is the feature used at node *n*, and Δ*I*_*n*_ is the information gain at node *n* where the feature *f* is used.

The feature importance profiles how the models leverage specific clinical indicators to predict fluid biomarkers. This knowledge can guide future efforts to streamline predictive feature sets (e.g., focusing on the most informative cognitive tests and demographics), and it provides confidence that the model is not relying on spurious or arbitrary noise correlations.

### Language Feature Extraction and Correlation Analysis

With surrogate biomarker values in hand for I-CONECT, we next examined their relationship to language-derived markers from the I-CONECT conversational data. All I-CONECT intervention sessions had been transcribed, and we extracted multiple quantitative linguistic features from these transcripts. We focused on four categories of language markers that have shown relevance to cognitive impairment in prior studies [19,20]: (1) Lexical diversity, reflecting the richness of vocabulary. We include ten lexical richness measures including Type-Token Ratio (TTR, Explanation - the ratio of unique words (types) to total words), Root TTR (Explanation - adjusts TTR by the square root of total tokens), and Maas TTR (Explanation - Sophisticated TTR variant accounting for text length). We calculated each measure of lexical diversity using Python’s lexical-diversity package; (2) Syntactic complexity, measuring the structural complexity of spoken sentences. We include twenty-three utterance measures, including nine frequency counts of the text structures (e.g., verb phrases and clauses) and fourteen syntactic complexity indices of the text (e.g., mean length of sentence and clauses per sentence). We calculated each measure of syntactic complexity using dependency parsers from Python Packages such as spaCy and Stanza. (3) LIWC categories (Linguistic Inquiry and Word Count), capturing psycholinguistic content such as pronoun usage, filler words, and sentiment-related words [1]. We include sixty-four LIWC features and extract those features using Python’s liwc-python package; and (4) Speech fluency metrics, including average response length (number of words per response or per turn) and frequency of pauses or hesitations. Supplementary Table 2 and Supplementary Table 3 in the appendix provide a detailed description for each lexical feature and syntactic feature.

For each I-CONECT participant, language features were aggregated over their conversation sessions (e.g. computing mean values per individual). We then computed correlation coefficients (ρ) between the predicted CSF biomarker values and each language feature, to assess whether individuals with greater estimated AD pathology show different language patterns.

### Code availability

The underlying code for this study is not publicly available but may be made available to qualified researchers on reasonable request from the corresponding author.

## Results

### Quality Assessment of Surrogate CSF Biomarker

All three CSF biomarker prediction models (LightGBM models) achieved reasonable accuracy in capturing variance in the biomarkers (for example, cross-validated R^2^ in the ∼0.5–0.6 range for Aβ42 prediction, indicating moderate explanatory power), as summarized in Table 2. Additionally, the corresponding prediction error plots are provided in Figure 2, where the values are log-scaled and normalized for better visualization.

**Table 2.** Performance of digital language features on regression tasks predicting surrogate CSF biomarkers (Aβ42, P-tau181, T-tau).

| $A\beta_{42}$ regression task | | P-tau181 regression task | | T-tau regression task | |
| --- | --- | --- | --- | --- | --- |
| MSE | $R^2$ | MSE | $R^2$ | MSE | $R^2$ |
| 0.711 | 0.545 | 0.871 | 0.279 | 0.659 | 0.540 |

**Figure 2.**
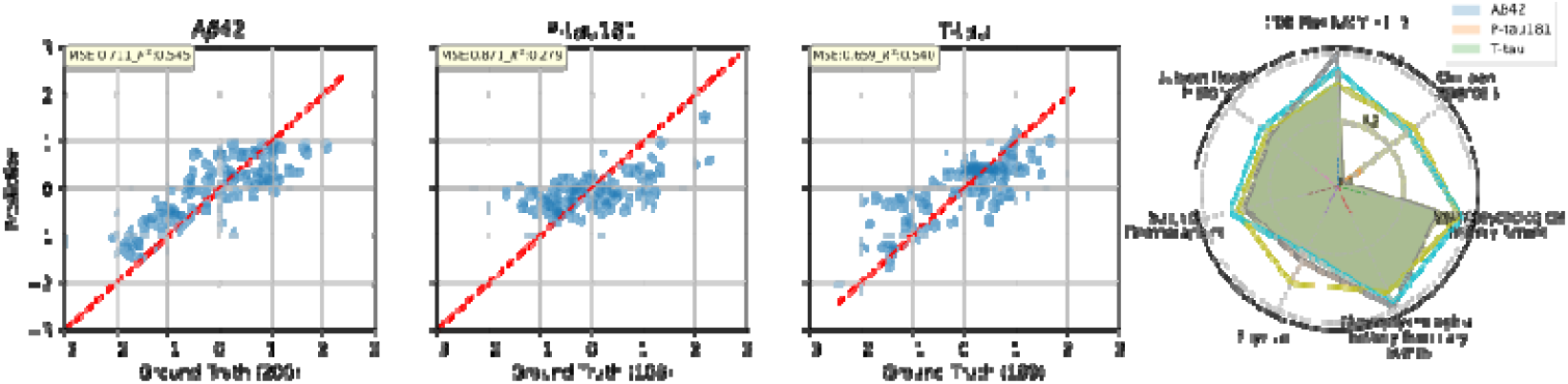
Prediction error plot for CSF Markers, Aβ42, P-tau181, and T-tau, regression tasks under 90:10 split for NACC data. And corresponding cumulative (log-scaled) feature importances across various feature categories.

Moreover, as shown in Table 3, the feature importance analysis of the trained models revealed intuitive and interpretable patterns: cognitive scores were among the strongest predictors of biomarker levels. For example, Clinical Dementia Rating (CDR) sub-items related to orientation and memory, as well as the clinician’s global assessment of cognitive status (e.g., ORIENT, MEMORY, and COGSTAT), were highly influential in the Aβ42 model. In contrast, neuropsychological test scores reflecting earlier stages of cognitive decline—preceding changes typically captured by CDR sub-items—ranked among the top predictors for P-tau181 (e.g., CRAFTDVR, CRAFTDTI, and MOCARECN). These findings are consistent with well-established associations between CSF biomarkers and cognition—namely, that lower CSF Aβ42 and higher tau levels are linked to poorer cognitive performance—and suggest that P-tau181 may be sensitive to earlier cognitive decline [21]. The dominance of a handful of features in each model (with a long tail of variables contributing smaller effects) implies that while the model casts a wide net (168 predictors), a core subset of measures carries most of the predictive power.

**Table 3.** Top 20 most important features for each regression task predicting surrogate CSF biomarkers (Aβ42, P-tau181, T-tau). Note: ORIENT_0 represents a categorical feature named ORIENT, with the category identified by the ID “0”.

| Aβ42 regression task | | P-tau181 regression task | | T-tau regression task | |
| --- | --- | --- | --- | --- | --- |
| Feature | Imp. | Feature | Imp. | Feature | Imp. |
| ORIENT_0.0 | 2704 | CRAFTDVR | 1458 | COGSTAT_0 | 2473 |
| COGSTAT_0 | 2287 | CRAFTDTI | 716 | CRAFTDTI | 2271 |
| CDRLANG_-4.0 | 1240 | COGSTAT_0 | 713 | CDRLANG_-4.0 | 1450 |
| COGSTAT_4 | 685 | MOCARECN | 556 | UDSBENTD | 625 |
| MEMORY_1.0 | 485 | DEMENTED_0 | 517 | CRAFTDVR | 454 |
| TRAILB | 247 | RACE_1 | 421 | DEMENTED_0 | 330 |
| MOCAORCT | 243 | UDSBENTD | 335 | PRIMLANG_1 | 280 |
| COGSTAT_3 | 237 | WEIGHT | 268 | RACE_1 | 235 |
| VEG | 228 | HRATE | 206 | HISPANIC_0 | 181 |
| MEMORY_0.5 | 227 | COMPORT_0.0 | 180 | RACE_2 | 165 |
| WEIGHT | 119 | MEMORY_0.0 | 146 | MINTSCNG | 123 |
| HISPANIC_0 | 119 | TRAILA | 131 | JUDGMENT_0.0 | 119 |
| MOCASER7 | 114 | PRIMLANG_2 | 113 | INCONTF_0 | 112 |
| EDUC | 111 | CDRLANG_-4.0 | 107 | TRAILBLI | 103 |
| ORIENT_0.5 | 96 | ORIENT_0.0 | 102 | PD_1 | 101 |
| TRAILBLI | 87 | BPSYS | 98 | MOCARECN | 99 |
| PRIMLANG_1 | 80 | UDSBENTC | 73 | MOCATOTS | 97 |
| UDSBENTC | 76 | HEIGHT | 72 | PRIMLANG_2 | 91 |
| TRAILALI | 60 | PD_1 | 64 | VISION_0 | 89 |
| MOCAREGI | 56 | SMOKYRS | 54 | ORIENT_0.0 | 85 |

### Visualization of Distribution Shift between NACC and I-CONECT data

The visualization of the distribution shift is illustrated in Figure 3. We observe that there are two main subgroups of subjects from the I-CONECT data that are out-of-distribution. The trained biomarker prediction models are less like to correctly predict surrogate CSF biomarker for those subjects. Hence, 30% of I-CONECT out-of-distribution cases is trimmed, the distribution overlap improved and the LightGBM models’ input features fell within a more reliable range. Notably, the language– biomarker correlations remained qualitatively similar after removing outliers, suggesting our findings are not driven by those extreme points but rather reflect a general trend.

**Figure 3.**
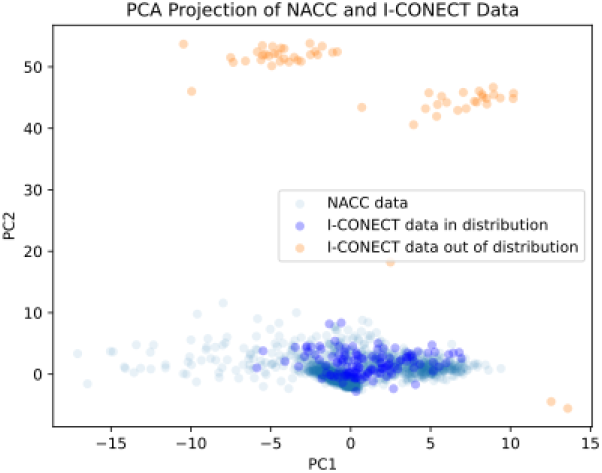
Distribution Shift Assessment between I-CONECT and NACC datasets over the 168-dimensional feature space: PCA project over NACC data and apply to I-CONECT data.

Finally, using the optimized LightGBM models, we then computed predicted CSF biomarker values for each I-CONECT participant by inputting their 168 common-feature data into the models. The output is a set of “surrogate” CSF biomarker estimates for the I-CONECT cohort, providing an approximate indication of each individual’s AD-related pathology burden (without requiring an invasive lumbar puncture). The distribution of CSF Marker prediction results regarding Aβ42, P-tau181, and T-tau is shown as Figure 4.

**Figure 4.**
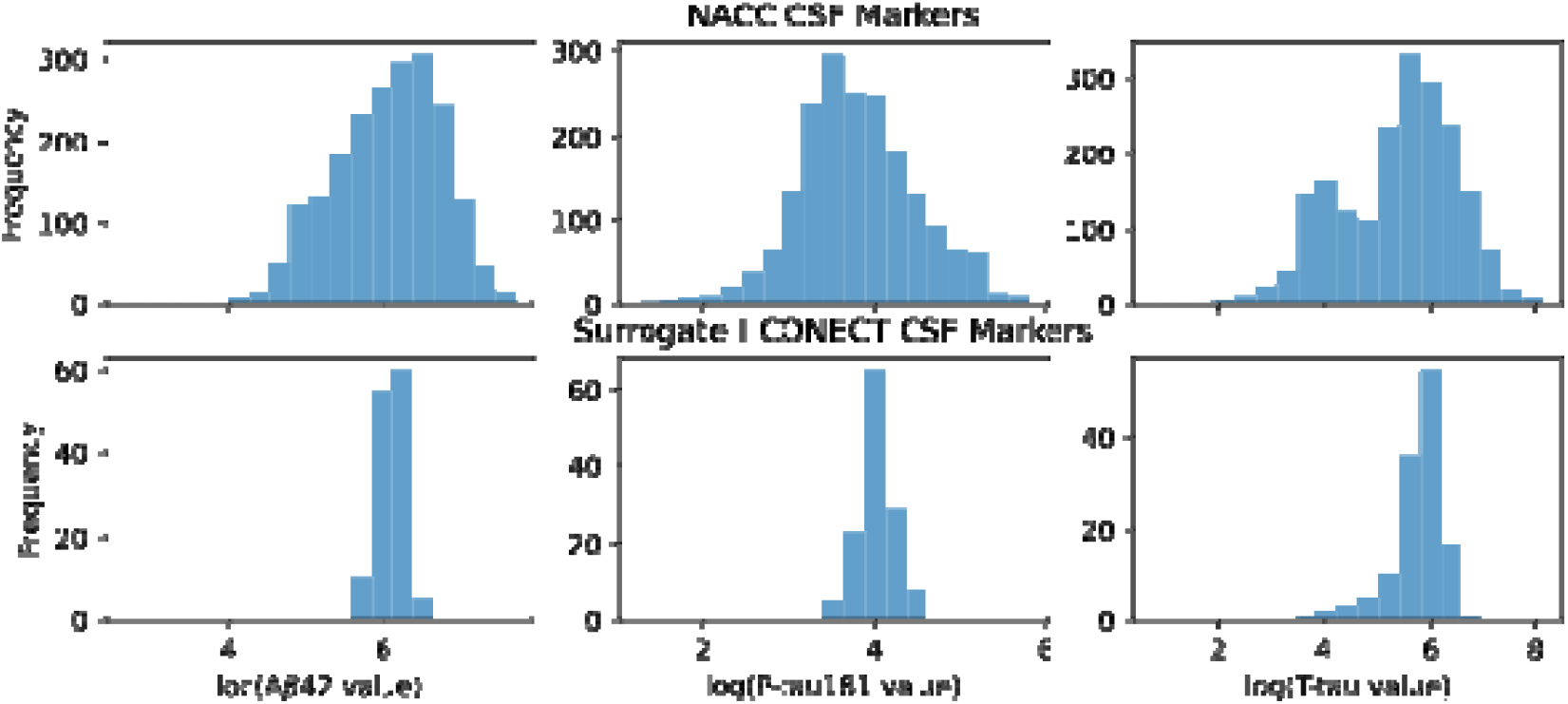
Predicted distribution of CSF biomarkers (Aβ42, P-tau181, and T-tau) for I-CONECT participants, compared with the reference distribution from NACC.

### Language Marker Correlation with Surrogate CSF Markers

We present the results of language marker correlation with each surrogate CSF marker in Table 4. Notably, we observed several significant correlations (two-tailed p<0.05 after correcting for multiple comparisons). For example, participants with lower predicted CSF Aβ42 (indicating higher amyloid burden) tended to produce shorter and less complex speech: their average response lengths were shorter (ρ ≈+0.3, indicating higher Aβ42 corresponds to longer responses) and their syntactic complexity was reduced (ρ≈+0.3), suggesting that greater amyloid pathology relates to simpler, more telegraphic speech. Similarly, higher surrogate tau levels were associated with lower syntactic complexity (ρ≈–0.3). These patterns are consistent with cognitive-linguistic changes seen in early Alzheimer’s and MCI – prior work has shown that people with cognitive impairment speak less fluently or with a restricted vocabulary [19,20]. Although the correlation magnitudes are modest, they suggest a meaningful link: participants estimated to have higher AD pathology tended to exhibit subtle language deficits.

**Table 4.** Language Marker Correlation with Surrogate CSF Markers: Aβ42, P-tau181, and T-tau. Features are Selected and Ordered with P value.

| Selected language markers vs. Aβ42 | | | |
| --- | --- | --- | --- |
| Feature | Category | Pearson correlation | P value <sup>2</sup> |
| T-unit | Syntactic | 0.514 | 0.0001 |
| Sentence | Syntactic | 0.502 | 0.0002 |
| Articles | LIWC | 0.421 | 0.0021 |
| Root TTR | Lexical | 0.404 | 0.0033 |
| Coordinate phrase | Syntactic | 0.394 | 0.0042 |
| Work | LIWC | 0.382 | 0.0056 |
| Achievement | LIWC | 0.377 | 0.0064 |
| Mean of response length | Speech Segment | 0.365 | 0.0084 |
| Prepositions | LIWC | 0.365 | 0.0085 |
| Adverbs | LIWC | 0.356 | 0.0104 |
| Selected language markers vs. P-tau181 |  |  |  |
| Ingestion | LIWC | -0.291 | 0.0383 |
| Hear | LIWC | -0.273 | 0.0529 |
| Perceptual processes | LIWC | -0.224 | 0.1135 |
| Mass TTR | Lexical | -0.215 | 0.1301 |
| Root TTR | Lexical | 0.206 | 0.1476 |
| Selected language markers vs. T-tau |  |  |  |
| Negations | LIWC | -0.330 | 0.0179 |
| Inhibition | LIWC | -0.300 | 0.0322 |
| Exclusive | LIWC | -0.290 | 0.0390 |
| Present tense | LIWC | -0.285 | 0.0429 |
| Verb phrase per T-unit | Syntactic | -0.273 | 0.0526 |

## Discussion

Our findings highlight a novel integration of traditional ADRC-derived data with emerging digital biomarkers of Alzheimer’s disease. By leveraging the extensive NACC clinical database to train machine learning models, we were able to project established CSF biomarkers onto an intervention cohort with linguistic markers, I-CONECT, that otherwise lacks fluid biomarker data. The result is a set of surrogate amyloid and tau indicators for each individual, which we show have meaningful correlations with language patterns in conversation. This approach illustrates a new paradigm for AD research: using computational methodologies to bridge biological measures and behavioral signals. To our knowledge, this is the first study to demonstrate that everyday speech features – captured through a home-based conversational intervention – relate to the pathological hallmark biomarkers of AD (albeit via predicted values). The ability to detect signals of amyloid or tau pathology from natural language is especially exciting in the context of early detection and intervention. Speech and language are increasingly recognized as sensitive indicators of cognitive change that can precede overt clinical symptoms [19]. Our results reinforce this, showing that participants with language profiles suggestive of impairment (e.g. low syntactic complexity) tended to be the same individuals whom our models flagged as having higher AD biomarker burden. In practical terms, this suggests that **non-invasive social conversation could serve as a proxy screening tool** for underlying brain changes – an approach that is far more scalable and accessible than widespread CSF or Positron Emission Tomography (PET) testing.

### Novelty and Contribution

This work lies at the intersection of dementia biomarker research and digital health technology. Previous studies have established CSF Aβ42, tau, and p-tau as highly specific markers of AD pathology [17], and more recently demonstrated that various speech/language features can distinguish MCI or early dementia from healthy aging [1,2,5]. We extend this literature by directly linking these two domains. The novelty is twofold: first, using surrogate biomarker modeling, we infer likely pathological status in individuals who have only clinical assessments; second, we connect those inferences to spontaneous speech metrics collected in a real-world social context. This method leverages the strength of the ADRC/NACC infrastructure – large, standardized datasets for model training – and applies it to unlock insights from a cutting-edge digital dataset. In line with the theme of the special issue, our study showcases how 40 years of ADRC data and 25 years of NACC enable new discoveries when combined with advanced digital tools and AI methods. The ADRC program’s commitment to uniform data collection made it possible to identify common data elements between NACC and I-CONECT. Without such data harmonization across studies, connecting fluid biomarkers to digital language outcomes would be extraordinarily difficult. Our success in finding meaningful correlations attests to the transformative impact of ADRC and NACC resources in the era of computational dementia research. We demonstrate that historical data can be repurposed in innovative ways: a model trained on clinic-based cohorts predicted biomarker levels in a completely independent sample, yielding results that are interpretable and clinically relevant. This underscores the ongoing relevancy of NACC’s repository – not only for traditional analyses, but as a foundation to calibrate novel digital biomarkers.

### Implications for Early Detection

Early and accurate detection of AD is a pressing goal, especially with the advent of disease-modifying therapies targeting amyloid and tau. Our approach could help identify individuals at risk before overt cognitive decline is evident, by using subtle linguistic changes as red flags. The finding that, for example, lower predicted Aβ42 correlates with simpler speech and smaller vocabulary suggests that amyloid pathology may manifest in everyday communication even in cognitively normal or mildly impaired older adults. If validated in further studies, one could envision a screening system where a short sample of a person’s conversations (perhaps obtained via a phone conversation or virtual assistant) is analyzed for certain linguistic markers, and in conjunction with a predictive model, estimates an “AD biomarker risk score.” Those above a certain risk threshold could be referred for confirmatory biomarker testing (such as CSF analysis or amyloid PET) or for early intervention strategies. This tiered approach would greatly widen the screening funnel using a cost-effective, non-invasive tool, aligning with precision health initiatives. Furthermore, because our surrogate biomarkers are on a continuous scale, they could be used to track changes within individuals. For instance, in a longitudinal setting, increasing deviation in a person’s predicted tau over time might signal accelerating neurodegeneration, prompting a clinical evaluation even if standard cognitive tests remain borderline. In the I-CONECT trial context, future analyses (as we proposed) will examine whether those with high predicted pathology benefit differently from the social engagement intervention or show divergent speech trajectories. This could inform personalized intervention strategies – perhaps tailoring cognitive engagement techniques for those with higher inferred pathology.

### Limitations and Future Directions

We acknowledge several limitations of this study. First, the surrogate biomarker estimates are inherently imperfect proxies. Given the secondary data analysis setting and the lack of alternative datasets with similarly semi-structured spoken language data (i.e., social conversations), external validation is currently not feasible. Thus, the clinical significance of the observed correlations between directly measured language features and machine-predicted surrogate biomarkers should be interpreted with caution, as the surrogate values have not been validated on an independent cohort. Especially, there is inevitable uncertainty in the predicted CSF values for any given individual. We present group-level correlations, which are more robust, but the approach would benefit from further validation against real biomarker measurements in the same individuals (e.g., if a subset of I-CONECT participants eventually provided blood or CSF samples for comparison). Second, domain shift between datasets is a challenge. The I-CONECT population, by design (e.g., socially isolated, older), is not fully representative of the broader ADRC cohorts that trained the model. We took steps to mitigate this (PCA-based outlier removal and cluster analysis to understand group differences) but acknowledge that these procedures may still introduce selection bias, potentially affecting the model’s accuracy and generalizability. More sophisticated domain adaptation techniques or recalibration of the model might improve accuracy. As more ADRC data become available – for example, if future UDS (Uniform Data Set) iterations incorporate digital voice recordings – it will be possible to retrain or fine-tune models on a more diverse sample, narrowing the gap between training and application domains. However, these recordings may still be limited to those obtained during neuropsychological testing, which offer constrained linguistic variability. Third, our language analysis thus far is cross-sectional (averaged over sessions), and the causal direction cannot be determined. It could be that underlying pathology leads to communication changes, but conversely, those with worse language abilities might be less cognitively stimulated and thus more prone to pathology – or an unknown third factor could influence both. Longitudinal analysis will help clarify the directionality: if those with high baseline surrogate biomarkers show greater decline in language measures over time, that strengthens the case that pathology drives the language changes. We are also aware that confounding factors like education, hearing loss, or even baseline depression could affect language use independent of AD pathology. Our regression models implicitly adjust for many such factors (since they’re included as features), but residual confounding is possible. In future work, we will explore multivariate models that include predicted biomarkers alongside demographic and health covariates to see if the biomarkers independently predict language outcomes. Despite these limitations, the present study provides a valuable proof-of-concept for linking conventional biomarkers with digital behavior data.

## Acknowledgements

NACC data are contributed by the NIA-funded ADRCs: P30 AG062429 (PI James Brewer, MD, PhD), P30 AG066468 (PI Oscar Lopez, MD), P30 AG062421 (PI Bradley Hyman, MD, PhD), P30 AG066509 (PI Thomas Grabowski, MD), P30 AG066514 (PI Mary Sano, PhD), P30 AG066530 (PI Helena Chui, MD), P30 AG066507 (PI Marilyn Albert, PhD), P30 AG066444 (PI David Holtzman, MD), P30 AG066518 (PI Lisa Silbert, MD, MCR), P30 AG066512 (PI Thomas Wisniewski, MD), P30 AG066462 (PI Scott Small, MD), P30 AG072979 (PI David Wolk, MD), P30 AG072972 (PI Charles DeCarli, MD), P30 AG072976 (PI Andrew Saykin, PsyD), P30 AG072975 (PI Julie A. Schneider, MD, MS), P30 AG072978 (PI Ann McKee, MD), P30 AG072977 (PI Robert Vassar, PhD), P30 AG066519 (PI Frank LaFerla, PhD), P30 AG062677 (PI Ronald Petersen, MD, PhD), P30 AG079280 (PI Jessica Langbaum, PhD), P30 AG062422 (PI Gil Rabinovici, MD), P30 AG066511 (PI Allan Levey, MD, PhD), P30 AG072946 (PI Linda Van Eldik, PhD), P30 AG062715 (PI Sanjay Asthana, MD, FRCP), P30 AG072973 (PI Russell Swerdlow, MD), P30 AG066506 (PI Glenn Smith, PhD, ABPP), P30 AG066508 (PI Stephen Strittmatter, MD, PhD), P30 AG066515 (PI Victor Henderson, MD, MS), P30 AG072947 (PI Suzanne Craft, PhD), P30 AG072931 (PI Henry Paulson, MD, PhD), P30 AG066546 (PI Sudha Seshadri, MD), P30 AG086401 (PI Erik Roberson, MD, PhD), P30 AG086404 (PI Gary Rosenberg, MD), P20 AG068082 (PI Angela Jefferson, PhD), P30 AG072958 (PI Heather Whitson, MD), P30 AG072959 (PI James Leverenz, MD). The I⍰CONECT project was supported by National Institutes of Health grants R01 AG051628 and R01 AG056102. The current project is supported by a National Institutes of Health grant R01AG072449.

## Data Availability

The data that support the findings of this study are available via NACC and I-CONECT but restrictions apply to the availability of these data, which were used under licence for the current study, and so are not publicly available.

## References

1. Asgari M, Kaye J, Dodge H. Predicting mild cognitive impairment from spontaneous spoken utterances. Alzheimers Dement (N Y) 2017;3:219–28.

2. Tang F, Chen J, Dodge HH, Zhou J. The joint effects of acoustic and linguistic markers for early identification of mild cognitive impairment. Front Digit Health 2021;3:702772.

3. Hoang B, Pang Y, Dodge HH, Zhou J. Subject harmonization of multi-modal digital markers: Improved detection of mild cognitive impairment using language and facial expression. Alzheimers Dement 2024;20. 10.1002/alz.094211.

4. Poor FF, Dodge HH, Mahoor MH. A multimodal cross-transformer-based model to predict mild cognitive impairment using speech, language and vision. Comput Biol Med 2024;182:109199.

5. Hoang B, Pang Y, Dodge H, Zhou J. Translingual language markers for cognitive assessment from spontaneous speech. Interspeech 2024, ISCA: ISCA; 2024, p. 977–81.

6. Cho S, Cousins KAQ, Shellikeri S, Ash S, Irwin DJ, Liberman MY, et al. Lexical and acoustic speech features relating to Alzheimer disease pathology. Neurology 2022;99:e313–22.

7. García-Gutiérrez F, Marquié M, Muñoz N, Alegret M, Cano A, de Rojas I, et al. Harnessing acoustic speech parameters to decipher amyloid status in individuals with mild cognitive impairment. Front Neurosci 2023;17:1221401.

8. van den Berg RL, de Boer C, Zwan MD, Jutten RJ, van Liere M, van de Glind M-CABJ, et al. Digital remote assessment of speech acoustics in cognitively unimpaired adults: feasibility, reliability and associations with amyloid pathology. Alzheimers Res Ther 2024;16:176.

9. Farzana S, Stoppa E, Leow A, Gollan T, Moore R, Salmon D, et al. SLaCAD: A spoken language corpus for early Alzheimer ‘s disease detection. In: Calzolari N, Kan M-Y, Hoste V, Lenci A, Sakti S, Xue N, editors. Proceedings of the 2024 Joint International Conference on Computational Linguistics, Language Resources and Evaluation (LREC-COLING 2024), Torino, Italia: ELRA and ICCL; 2024, p. 14877–97.

10. Chou C-J, Chang C-T, Chang Y-N, Lee C-Y, Chuang Y-F, Chiu Y-L, et al. Screening for early Alzheimer’s disease: enhancing diagnosis with linguistic features and biomarkers. Front Aging Neurosci 2024;16:1451326.

11. Dodge HH, Yu K, Wu C-Y, Pruitt PJ, Asgari M, Kaye JA, et al. Internet-based conversational engagement randomized controlled clinical trial (I-CONECT) among socially isolated adults 75+ years old with normal cognition or mild cognitive impairment: Topline results. Gerontologist 2024;64. 10.1093/geront/gnad147.

12. ViViT: Multi-branch Classifier-ViViT to detect Mild Cognitive Impairment in older adults using facial videos. n.d.

13. Liu G, Xue Z, Zhan L, Dodge HH, Zhou J. Detection of mild cognitive impairment from language markers with crossmodal augmentation. Pac Symp Biocomput 2023;28:7–18.

14. Pourramezan Fard A, Mahoor MH, Alsuhaibani M, Dodge HH. Linguistic-based Mild Cognitive Impairment detection using Informative Loss. Comput Biol Med 2024;176:108606.

15. Besser L, Kukull W, Knopman DS, Chui H, Galasko D, Weintraub S, et al. Version 3 of the national Alzheimer’s coordinating center’s Uniform Data Set. Alzheimer Dis Assoc Disord 2018;32:351–8.

16. White MT, Shaw LM, Xie SX, Alzheimer’s Disease Neuroimaging Initiative, National Alzheimer’s Coordinating Center. Evaluation of cerebrospinal fluid assay variability in Alzheimer’s disease. J Alzheimers Dis 2016;51:463–70.

17. Humpel C. Identifying and validating biomarkers for Alzheimer’s disease. Trends Biotechnol 2011;29:26–32.

18. Hajihosseinlou M, Maghsoudi A, Ghezelbash R. A novel scheme for mapping of MVT-type Pb–Zn prospectivity: LightGBM, a highly efficient gradient boosting decision tree machine learning algorithm. Nat Resour Res 2023;32:2417–38.

19. Lima MR, Capstick A, Geranmayeh F, Nilforooshan R, Matarić M, Vaidyanathan R, et al. Evaluating spoken language as a biomarker for automated screening of cognitive impairment. ArXiv [CsLG] 2025.

20. Boland AK, Jensen A, Davidson PSR, Taler V. Linguistic markers of story recall can help differentiate mild cognitive impairment from normal aging. Language and Health 2024;2:100030.

21. Skillbäck T, Farahmand BY, Rosén C, Mattsson N, Nägga K, Kilander L, et al. Cerebrospinal fluid tau and amyloid-β1-42 in patients with dementia. Brain 2015;138:2716–31.

