## Supplemental tables for "Bridging Language Markers and Pathology: Correlations Between Digital Speech Measures and Surrogate CSF Biomarkers in Alzheimer’s Disease"

### Appendix

Supplementary Table 1 Handbook of the 168 common variables collected in both datasets: I-CONECT trial and the NACC database.

| Data Dictionary Codebook | | |
| --- | --- | --- |
| Variable | Form | Descriptor |
| SEX | Subject Demographics | Subject's sex |
| HISPANIC | Subject Demographics | Hispanic/Latino ethnicity |
| HISPOR | Subject Demographics | Hispanic origins |
| HISPORX | Subject Demographics | Hispanic origins, other - specify |
| RACE | Subject Demographics | Race |
| RACEX | Subject Demographics | Race, other - specify |
| RACESEC | Subject Demographics | Second race |
| RACESECX | Subject Demographics | Second race, other - specify |
| RACETER | Subject Demographics | Third race |
| RACETERX | Subject Demographics | Third race, other - specify |
| PRIMLANG | Subject Demographics | Primary language |
| PRIMLANX | Subject Demographics | Primary language, other- specify |
| EDUC | Subject Demographics | Years of education |
| MARISTAT | Subject Demographics | Marital status |
| INDEPEND | Subject Demographics | Level of independence |
| RESIDENC | Subject Demographics | Type of residence |
| HANDED | Subject Demographics | Is the subject left- or right-handed? |
| TOBAC30 | Subject Health History | Smoked cigarettes in last 30 days |
| TOBAC100 | Subject Health History | Smoked more than 100 cigarettes in life |
| SMOKYRS | Subject Health History | Total years smoked cigarettes |
| PACKSPER | Subject Health History | Average number of packs smoked per day |
| QUITSMOK | Subject Health History | If the subject quit smoking, age at which he/she last smoked (i.e., quit) |
| ALCOCCAS | Subject Health History | In the past three months, has the subject consumed any alcohol? |
| ALCFREQ | Subject Health History | During the past three months, how often did the subject have at least one drink of any alcoholic beverage such as wine, beer, malt liquor, or spirits? |
| CVHATT | Subject Health History | Heart attack/cardiac arrest |
| HATTMULT | Subject Health History | More than one heart attack/cardiac arrest? |
| HATTYEAR | Subject Health History | Year of most recent heart attack |
| CVAFIB | Subject Health History | Atrial fibrillation |
| CVANGIO | Subject Health History | Angioplasty/endarterectomy/stent |
| CVBYPASS | Subject Health History | Cardiac bypass procedure |
| CVPACDEF | Subject Health History | Pacemaker and/or defibrillator |
| CVCHF | Subject Health History | Congestive heart failure |
| CVANGINA | Subject Health History | Angina |
| CVHVALVE | Subject Health History | Heart valve replacement or repair |
| CVOTHR | Subject Health History | Other cardiovascular disease |
| CVOTHRX | Subject Health History | Specification for other cardiovascular disease |
| CBSTROKE | Subject Health History | Stroke |
| STROKMUL | Subject Health History | More than one stroke reported as of the Initial Visit |
| CBTIA | Subject Health History | Transient ischemic attack (TIA) |
| TIAMULT | Subject Health History | More than one TIA reported as of the Initial Visit |
| PD | Subject Health History | Parkinson's disease (PD) |
| PDYR | Subject Health History | Year of PD diagnosis |
| PDOTHR | Subject Health History | Other parkinsonian disorder |
| PDOTHRYR | Subject Health History | Year of parkinsonian disorder diagnosis |
| SEIZURES | Subject Health History | Seizures |
| TBI | Subject Health History | Traumatic brain injury (TbI) |
| TBIBRIEF | Subject Health History | Traumatic brain injury (TbI) with brief loss of consciousness |
| TBIEXTEN | Subject Health History | TbI with extended loss of consciousness - 5 minutes of longer |
| TBIWOLOS | Subject Health History | TbI without loss of consciousness - as might result from military detonations or sports injury |
| TBIYEAR | Subject Health History | Year of most recent TbI |
| DIABETES | Subject Health History | Diabetes |
| DIABTYPE | Subject Health History | If Recent/active or Remote/inactive diabetes, which type? |
| HYPERTEN | Subject Health History | Hypertension |
| HYPERCHO | Subject Health History | Hypercholesterolemia |
| B12DEF | Subject Health History | Vitamin b12 deficiency |
| THYROID | Subject Health History | Thyroid disease |
| ARTHRIT | Subject Health History | Arthritis |
| ARTHTYPE | Subject Health History | Type of arthritis |
| ARTHTYPX | Subject Health History | Other arthritis (specify) |
| ARTHUPEX | Subject Health History | Arthritis, region affected - upper extremity |
| ARTHLOEX | Subject Health History | Arthritis, region affected - lower extremity |
| ARTHSPIN | Subject Health History | Arthritis, region affected - spine |
| ARTHUNK | Subject Health History | Region affected - unknown |
| INCONTU | Subject Health History | Incontinence - urinary |
| INCONTF | Subject Health History | Incontinence - bowel |
| APNEA | Subject Health History | Sleep apnea history reported at Initial Visit |
| RBD | Subject Health History | REM sleep behavior disorder (RbD) history reported at Initial Visit |
| INSOMN | Subject Health History | Hyposomnia/insomnia history reported at Initial Visit |
| OTHSLEEP | Subject Health History | Other sleep disorder history reported at Initial Visit |
| OTHSLEEX | Subject Health History | Other sleep disorder (specify) |
| ALCOHOL | Subject Health History | Alcohol abuse - clinically significant occurring over a 12-month period manifested in one of the following areas: work, driving, legal, or social |
| ABUSOTHR | Subject Health History | Other abused substances - clinically significant impairment occurring over a 12-month period manifested in one of the following areas: work, driving, legal, or social |
| ABUSX | Subject Health History | If reported other abused substances, specify abused substance(s) |
| BIPOLAR | Subject Health History | bipolar disorder |
| SCHIZ | Subject Health History | Schizophrenia |
| DEP2YRS | Subject Health History | Active depression in the last two years |
| DEPOTHR | Subject Health History | Depression episodes more than two years ago |
| ANXIETY | Subject Health History | Anxiety |
| OCD | Subject Health History | Obsessive-compulsive disorder (OCD) |
| NPSYDEV | Subject Health History | Developmental neuropsychiatric disorders (e.g., autism spectrum disorder [ASD], attention-deficit hyperactivity disorder [ADHD], dyslexia) |
| PSYCDIS | Subject Health History | Other psychiatric disorder |
| PSYCDISX | Subject Health History | If recent/active or remote/inactive psychiatric disorder, specify disorder |
| HEIGHT | Physical | Subject's height (inches) |
| WEIGHT | Physical | Subject's weight (lbs) |
| BPSYS | Physical | Subject blood pressure (sitting), systolic |
| BPDIAS | Physical | Subject blood pressure (sitting), diastolic |
| HRATE | Physical | Subject resting heart rate (pulse) |
| VISION | Physical | Without corrective lenses, is the subject's vision functionally normal? |
| VISCORR | Physical | Does the subject usually wear corrective lenses? |
| VISWCORR | Physical | If the subject usually wears corrective lenses, is the subject's vision functionally normal with corrective lenses? |
| HEARING | Physical | Without a hearing aid(s), is the subject's hearing functionally normal? |
| HEARAID | Physical | Does the subject usually wear a hearing aid(s)? |
| HEARWAID | Physical | If the subject usually wears a hearing aid(s), is the subject's hearing functionally normal with a hearing aid(s)? |
| MEMORY | CDR Plus NACC FTLD | Memory |
| ORIENT | CDR Plus NACC FTLD | Orientation |
| JUDGMENT | CDR Plus NACC FTLD | Judgment and problem-solving |
| COMMUN | CDR Plus NACC FTLD | Community affairs |
| HOMEHOBB | CDR Plus NACC FTLD | Home and hobbies |
| PERSCARE | CDR Plus NACC FTLD | Personal care |
| CDRSUM | CDR Plus NACC FTLD | CDR sum of boxes |
| CDRGLOB | CDR Plus NACC FTLD | Global CDR |
| COMPORT | CDR Plus NACC FTLD | behavior, comportment, and personality |
| CDRLANG | CDR Plus NACC FTLD | Language |
| UDSBENTC | Neuropsychological battery Summary Scores | Total score for copy of benson figure |
| UDSBENTD | Neuropsychological battery Summary Scores | Total score for 10- to 15-minute delayed drawing of benson figure |
| UDSBENRS | Neuropsychological battery Summary Scores | Recognized original stimulus from among four options |
| ANIMALS | Neuropsychological battery Summary Scores | Animals - Total number of animals named in 60 seconds |
| VEG | Neuropsychological battery Summary Scores | Vegetable - Total number of vegetables named in 60 seconds |
| TRAILA | Neuropsychological battery Summary Scores | Trail Making Test Part A - Total number of seconds to complete |
| TRAILARR | Neuropsychological battery Summary Scores | Part A - Number of commission errors |
| TRAILALI | Neuropsychological battery Summary Scores | Part A - Number of correct lines |
| TRAILB | Neuropsychological battery Summary Scores | Trail Making Test Part b - Total number of seconds to complete |
| TRAILBRR | Neuropsychological battery Summary Scores | Part b - Number of commission errors |
| TRAILBLI | Neuropsychological battery Summary Scores | Part b - Number of correct lines |
| UDSVERFC | Neuropsychological battery Summary Scores | Number of correct F-words generated in 1 minute |
| UDSVERFN | Neuropsychological battery Summary Scores | Number of F-words repeated in 1 minute |
| UDSVERNF | Neuropsychological battery Summary Scores | Number of non-F-words and rule violation errors in 1 minute |
| UDSVERLC | Neuropsychological battery Summary Scores | Number of correct L-words generated in 1 minute |
| UDSVERLR | Neuropsychological battery Summary Scores | Number of L-words repeated in 1 minute |
| UDSVERLN | Neuropsychological battery Summary Scores | Number of non-L-words and rule violation errors in 1 minute |
| UDSVERTN | Neuropsychological battery Summary Scores | Total number of correct F-words and L-words |
| UDSVERTE | Neuropsychological battery Summary Scores | Total number of F-word and L-word repetition errors |
| UDSVERTI | Neuropsychological battery Summary Scores | Total number of non-F/L-words and rule violation errors |
| COGSTAT | Neuropsychological battery Summary Scores | Per clinician, based on the neuropsychological examination, the subject's cognitive status is deemed |
| MOCAVIS | Neuropsychological battery Scores | Subject was unable to complete one or more sections due to visual impairment |
| MOCAHEAR | Neuropsychological battery Scores | Subject was unable to complete one or more sections due to hearing impairment |
| MOCATOTS | Neuropsychological battery Scores | MoCA Total Raw Score - uncorrected |
| MOCATRAI | Neuropsychological battery Scores | MoCA: Visuospatial/executive - Trails |
| MOCACUBE | Neuropsychological battery Scores | MoCA: Visuospatial/executive - Cube |
| MOCACLOC | Neuropsychological battery Scores | MoCA: Visuospatial/executive - Clock contour |
| MOCACLON | Neuropsychological battery Scores | MoCA: Visuospatial/executive - Clock numbers |
| MOCACLOH | Neuropsychological battery Scores | MoCA: Visuospatial/executive - Clock hands |
| MOCANAMI | Neuropsychological battery Scores | MoCA: Language - Naming |
| MOCAREGI | Neuropsychological battery Scores | MoCA: Memory - Registration (two trials) |
| MOCADIGI | Neuropsychological battery Scores | MoCA: Attention - Digits |
| MOCALETT | Neuropsychological battery Scores | MoCA: Attention - Letter A |
| MOCASER7 | Neuropsychological battery Scores | MoCA: Attention - Serial 7s |
| MOCAREPE | Neuropsychological battery Scores | MoCA: Language - Repetition |
| MOCAFLUE | Neuropsychological battery Scores | MoCA: Language - Fluency |
| MOCAABST | Neuropsychological battery Scores | MoCA: Abstraction |
| MOCARECN | Neuropsychological battery Scores | MoCA: Delayed recall - No cue |
| MOCARECC | Neuropsychological battery Scores | MoCA: Delayed recall - Category cue |
| MOCARECR | Neuropsychological battery Scores | MoCA: Delayed recall - Recognition |
| MOCAORDT | Neuropsychological battery Scores | MoCA: Orientation - Date |
| MOCAORMO | Neuropsychological battery Scores | MoCA: Orientation - Month |
| MOCAORYR | Neuropsychological battery Scores | MoCA: Orientation - Year |
| MOCAORDY | Neuropsychological battery Scores | MoCA: Orientation - Day |
| MOCAORPL | Neuropsychological battery Scores | MoCA: Orientation - Place |
| MOCAORCT | Neuropsychological battery Scores | MoCA: Orientation - City |
| CRAFTVRS | Neuropsychological battery Scores | Craft Story 21 Recall (Immediate) - Total story units recalled, verbatim scoring |
| CRAFTURS | Neuropsychological battery Scores | Craft Story 21 Recall (Immediate) - Total story units recalled, paraphrase scoring |
| DIGFORCT | Neuropsychological battery Scores | Number Span Test: Forward - Number of correct trials |
| DIGFORSL | Neuropsychological battery Scores | Number Span Test: Forward - Longest span forward |
| DIGBACCT | Neuropsychological battery Scores | Number Span Test: backward - Number of correct trials |
| DIGBACLS | Neuropsychological battery Scores | Number Span Test: backward - Longest span backward |
| CRAFTDVR | Neuropsychological battery Scores | Craft Story 21 Recall (Delayed) - Total story units recalled, verbatim scoring |
| CRAFTDRE | Neuropsychological battery Scores | Craft Story 21 Recall (Delayed) - Total story units recalled, paraphrase scoring |
| CRAFTDTI | Neuropsychological battery Scores | Craft Story 21 Recall (Delayed) - Delay time |
| CRAFTCUE | Neuropsychological battery Scores | Craft Story 21 Recall (Delayed) - Cue (boy) needed |
| MINTTOTS | Neuropsychological battery Scores | Multilingual Naming Test (MINT) - Total score |
| MINTTOTW | Neuropsychological battery Scores | Multilingual Naming Test (MINT) - Total correct without semantic cue |
| MINTSCNG | Neuropsychological battery Scores | Multilingual Naming Test (MINT) - Semantic cues: Number given |
| MINTSCNC | Neuropsychological battery Scores | Multilingual Naming Test (MINT) - Semantic cues: Number correct with cue |
| MINTPCNG | Neuropsychological battery Scores | Multilingual Naming Test (MINT) - Phonemic cues: Number given |
| MINTPCNC | Neuropsychological battery Scores | Multilingual Naming Test (MINT) - Phonemic cues: Number correct with cue |
| NORMCOG | Clinician Diagnosis | Normal cognition and behavior |
| DEMENTED | Clinician Diagnosis | Met criteria for dementia |
| IMPNOMCI | Clinician Diagnosis | Cognitively impaired, not MCI |

Supplementary Table 2 Summary of lexical features: measuring a different aspect of lexical diversity in a given text.

| Lexical features | |
| --- | --- |
| Feature | Formula, Explanation, and Interpretation |
| Type-Token Ratio (TTR) | Formula - V/N, Explanation - Ratio of unique words (types) to total words (tokens), Interpretation - Higher values indicate greater vocabulary diversity, but sensitive to text length. |
| Root TTR | Formula – V/sqrt(N), Explanation - Adjusts TTR by the square root of total tokens, Interpretation - Reduces TTR sensitivity to text length. |
| Logarithmic TTR (Log TTR) | Formula - log⁡V/log⁡N, Explanation - Normalizes TTR using logarithms, Interpretation - Further reduces sensitivity to text length. |
| Maas TTR | Formula - (log⁡N−log⁡V)/(logN)^2, Explanation - Sophisticated TTR variant accounting for text length, Interpretation - Lower values indicate higher diversity. |
| Mean Segmental TTR | Formula - Average TTR over equal-sized text segments, Explanation - Reduces the impact of text length, Interpretation - More stable TTR measurement. |
| Moving Average TTR | Formula - Average TTR over overlapping text windows, Explanation - Smooths TTR across the text, Interpretation - Provides a consistent measure of diversity. |
| Hypergeometric Distribution Diversity (HDD) | Formula - Based on hypergeometric distribution, Explanation - Estimates diversity probabilistically, Interpretation - Less sensitive to text length. |
| Measure of Textual Lexical Diversity (MTLD) | Formula - Average segment length required to maintain a constant TTR threshold, Explanation - Measures consistency of lexical diversity, Interpretation - Higher values indicate greater diversity. |
| MTLD MA Wrap | Formula - MTLD calculated in both forward and backward directions, Explanation - Provides a more robust MTLD value, Interpretation - More stable than standard MTLD. |
| MTLD MA Bi-directional | Formula - MTLD calculated bidirectionally, Explanation - Balances MTLD from two directions, Interpretation - Provides a balanced view of MTLD. |

Supplementary Table 3 Summary of syntactic features.

| Syntactic features | |
| --- | --- |
| nine frequency counting of the text structures | words (W), sentences (S), verb phrases (VP), clauses (C), terminable-units (T, T-units), dependent clauses (DC), complex T-units (CT), coordinate phrases (CP), and complex nominals (CN) |
| fourteen syntactic complexity indices of the text | mean length of sentence (MLS), mean length of T-unit (MLT), mean length of clause (MLC), clauses per sentence (C/S), verb phrases per T-unit (VP/T), clauses per T-unit (C/T), dependent clauses per clause (DC/C), dependent clauses per T-unit (DC/T), T-units per sentence (T/S), complex T-unit ratio (CT/T), coordinate phrases per T-unit (CP/T), coordinate phrases per clause (CP/C), complex nominals per T-unit (CN/T), and complex nominals per clause (CN/C) |
